# Personalized transcranial alternating current stimulation improves sleep quality: Initial Findings

**DOI:** 10.1101/2022.09.26.509537

**Authors:** V. Ayanampudi, V. Kumar, A. Krishnan, M.P. Walker, R.B. Ivry, R.T. Knight, R. Gurumoorthy

## Abstract

Insufficient sleep is a major health issue. Inadequate sleep is associated with an array of poor health outcomes, including cardiovascular disease, diabetes, obesity, certain forms of cancer, Alzheimer’s disease, depression, anxiety, and suicidality. Given concerns with typical sedative hypnotic drugs for treating sleep difficulties, there is a compelling need for added alternative interventions. Here, we report results of a non-invasive electrical brain stimulation approach to optimizing sleep involving transcranial alternating current stimulation (tACS).

A total of 25 participants (mean age: 46.3, S.D. ±12.4, 15 females) were recruited for a null-stimulation controlled (Control condition), within subjects, randomized crossed design, that included two variants of an active condition involving 15 minutes pre-sleep tACS stimulation. To evaluate the impact on sleep quality, the two active tACS stimulation conditions were designed to modulate sleep-dependent neural activity in the theta/alpha frequency bands, with both stimulation types applied to all subjects in separate sessions. The first tACS condition used a fixed stimulation pattern across all participants, a pattern composed of stimulation at 5Hz and 10Hz. The second tACS condition used a personalized stimulation approach with the stimulation frequencies determined by each individual’s peak EEG frequencies in the 4-6Hz and 9-11Hz bands.

Personalized tACS stimulation increased sleep quantity (duration) by 22 minutes compared to a Control condition (p=.04), and 19 minutes compared to Fixed tACS stimulation (p=.03). Fixed stimulation did not significantly increase sleep duration compared to Control (mean: 3 minutes; p=0.75). For sleep onset, the Personalized tACS stimulation resulted in reducing the onset by 28% compared to the Fixed tACS stimulation (6 minutes faster, p=.02). For a Poor Sleep sub-group (n=13) categorized with Clinical Insomnia and with a high insomnia severity, Personalized tACS stimulation improved sleep duration by 33 minutes compared to Fixed stimulation (p=0.02), and 30 minutes compared to Control condition (p<0.1).

Together, these results suggest that Personalized stimulation improves sleep quantity and time taken to fall asleep relative to Control and Fixed stimulation providing motivation for larger-scale trials for Personalized tACS as a sleep therapeutic, including for those with insomnia.

## Introduction

Sleep is essential for human physical and mental health ^[1]^. This encompasses the homeostatic regulation of very major physiological systems within the body (e.g. cardiovascular, immune, endocrine, metabolic), as well as cognitive and emotional operations of the brain. Nevertheless, one in three adults average less than the CDC’s recommendation of at least seven hours ^[4]^. Moreover, the life-time incidence of insomnia in general population is estimated to be over 30% ^[25][4]^.

Conventional sleeping pills are widely used as therapeutic interventions for sleep difficulties. However, such sedative hypnotics have significant side effects, lack long-term efficacy, and may fail to restore normative sleep ^[2]^. Indeed, based on concerns of safety, efficacy, and side effects, sleeping pills are no longer recommended as a first line treatment approach for those with sleeping difficulties _[2]_.

These limitations have motivated investigations of non-pharmacological approaches to enhance sleep hygiene. One approach involves non-invasive brain stimulation whereby different forms of exogenous stimulation are used to modify sleep. To date, examples of such non-invasive methods include acoustic, magnetic, and electrical stimulation approaches targeting sleep onset, sleep quality and/or sleep duration ^[3]^.

The primary electrical stimulation methods are transcranial direct current stimulation (tDCS), and transcranial alternating current stimulation (tACS). Both involve the application of a low-intensity electrical current (1 – 2 mA) to the scalp. The main difference is that tDCS involves a fixed current while tACS uses an alternating current. tACS affords the advantage of creating flexible stimulation waveforms that can be manipulated to induce modulation of neural activity in targeted frequency bands.

Electrical brain stimulation has an appealing safety profile, with no adverse side effects ^[13]^. Indeed, data collected from 33,000 sessions and over 1,000 subjects who received repeated tDCS sessions, indicates that tDCS is safe to use in human subjects for repeated use. Furthermore, tDCS currents of up to 2mA have been shown to be safely tolerated ^[14] [15]^. tACS has similarly been reported to be safe without any adverse side effects ^[17]^ and has the added advantage of inducing less stimulation sensation on the scalp for the user, relative to tDCS ^[16]^.

Brain oscillatory patterns are grouped into different region-specific frequency bands critical for different brain functions ^[9] [18][19][20]^. Regarding sleep, scalp EEG and intracranial EEG investigations have provided evidence that electrical signatures from the frontal lobes are useful biomarkers of sleep onset and sleep maintenance ^[5][24]^. Mechanistically, it has been hypothesized that frontal lobe neurons provide “top-down” control of cortico-thalamic feedback loops involved in sleep-wake regulation ^[6]^.

Motivated by these observations, tDCS applied over the prefrontal cortex has been shown to beneficially modulate sleep ^[6]^. For example, application of 5Hz stimulation for 10 minutes to the frontal cortex of subjects increased subjective sleepiness and enhances slow-frequency EEG activity in the 0.5-4.75Hz range ^[7]^. Furthermore, bilateral 5Hz tACS stimulation over fronto-temporal areas decreases sleep onset ^[30]^ and increases slow wave activity in the first half of NREM cycle relative to sham ^[8]^. Further studies have shown the association of alpha band activity to drowsiness and transition from wakefulness to sleepiness ^[31]^.

To date, studies of the effect of tACS on sleep have used the same stimulation protocol for all participants, with the focusing on frequencies in the 0.5Hz to 16Hz range ^[32]^. However, EEG peak frequencies within specific bands show significant inter-individual differences ^[33][34]^. This has motivated the recent exploration of personalized stimulation protocols that are tailored to each participant’s innate frequency profile ^[26]^. Indeed customized alpha frequency stimulation during wakefulness, close to an individual’s internal alpha frequency improves the efficacy of stimulation entrainment, measures of underlying neural plasticity, and consequentially superior memory function _[27][12[28]_.

Based on these observations, the present study sought to test the hypothesis that tACS stimulation patterns prior to sleep was effective in improving sleep, and furthermore, tested whether personalized frequency closed loop stimulation was superior to fixed frequency stimulation.

Accordingly, the study employed two tACS approaches, one using a fixed stimulation protocol and the other using an individually tailored stimulation protocol. Participants were tested in their home using a custom designed headband that provided bilateral stimulation through two frontal electrodes. Using an app-based control system, stimulation was self-administered for 15 min just before the participant went to sleep. Two specific predictions were tested: 1) both types of tACS stimulation would improve total sleep duration and the time taken to fall asleep relative to a null-stimulation control condition, and 2) personalized tACS stimulation would provide superior sleep improvement on both metrics relative to fixed tACS stimulation.

## Methods

### Overview

Participants (detailed described, below) were tested over multiple sessions in their homes, self-administering the stimulation (Fixed tACS, Personalized tACS, or control) using a custom-built stimulator composed of a headband that contained two electrodes positioned over the frontal lobes. The participants used a customized phone app that randomly determined the stimulation mode for each of the alternating weeks. Sleep onset and duration were measured with a wearable tracker (Fitbit watch). The participants put the headband and tracker on when they were ready to go to sleep and used the phone app to start the stimulation.

### Participants and Protocols

A total of 25 participants were tested in a repeated-measures, cross-over design (mean age: 46.3, with the age ranging between 19-60, 10 male and 15 female – **Figure 1**). Participants completed the Insomnia Severity Index questionnaire prior to their first session using an online form (**Figure 2**). There was substantive representation of the ISI categories across the participants, with 28% categorized with no insomnia, 20% with subthreshold insomnia, 36% with clinical insomnia, and 16% with severe insomnia.

**Figure 1:**
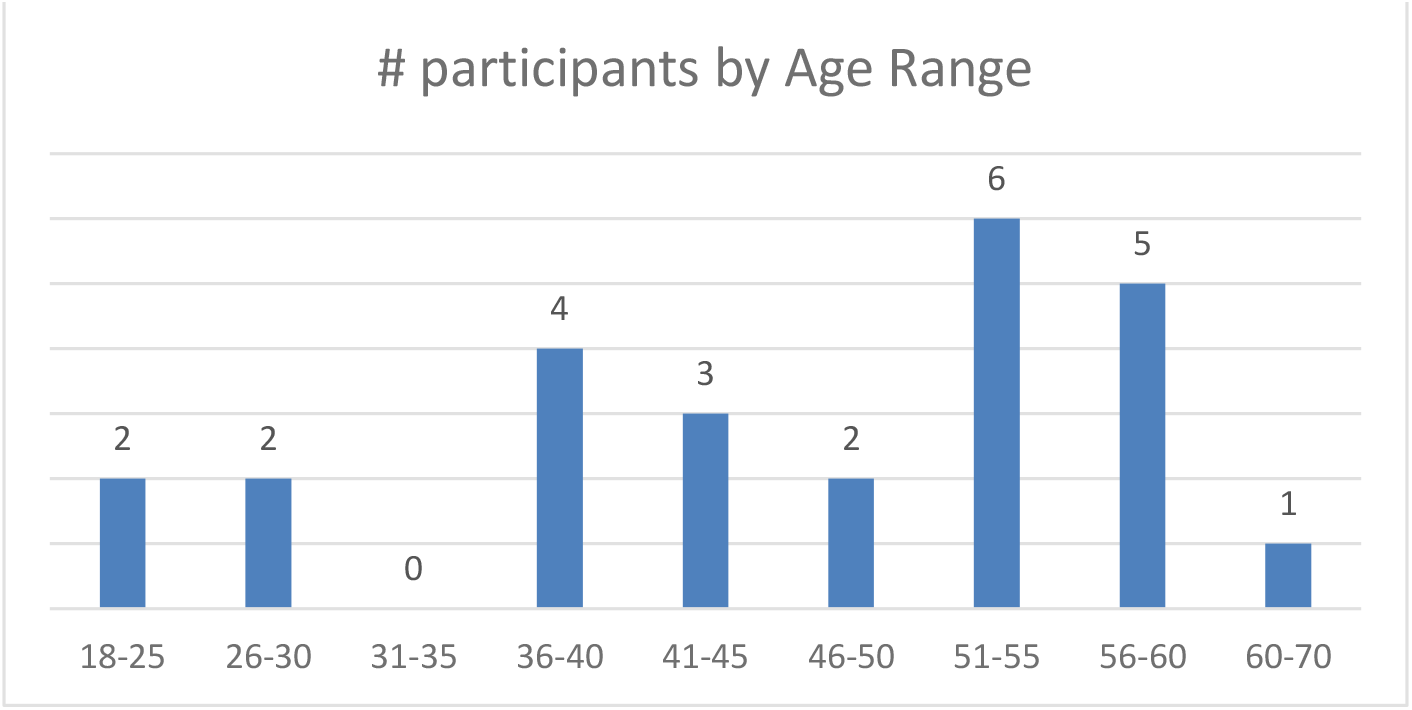
Age distribution of the participants

**Figure 2:**
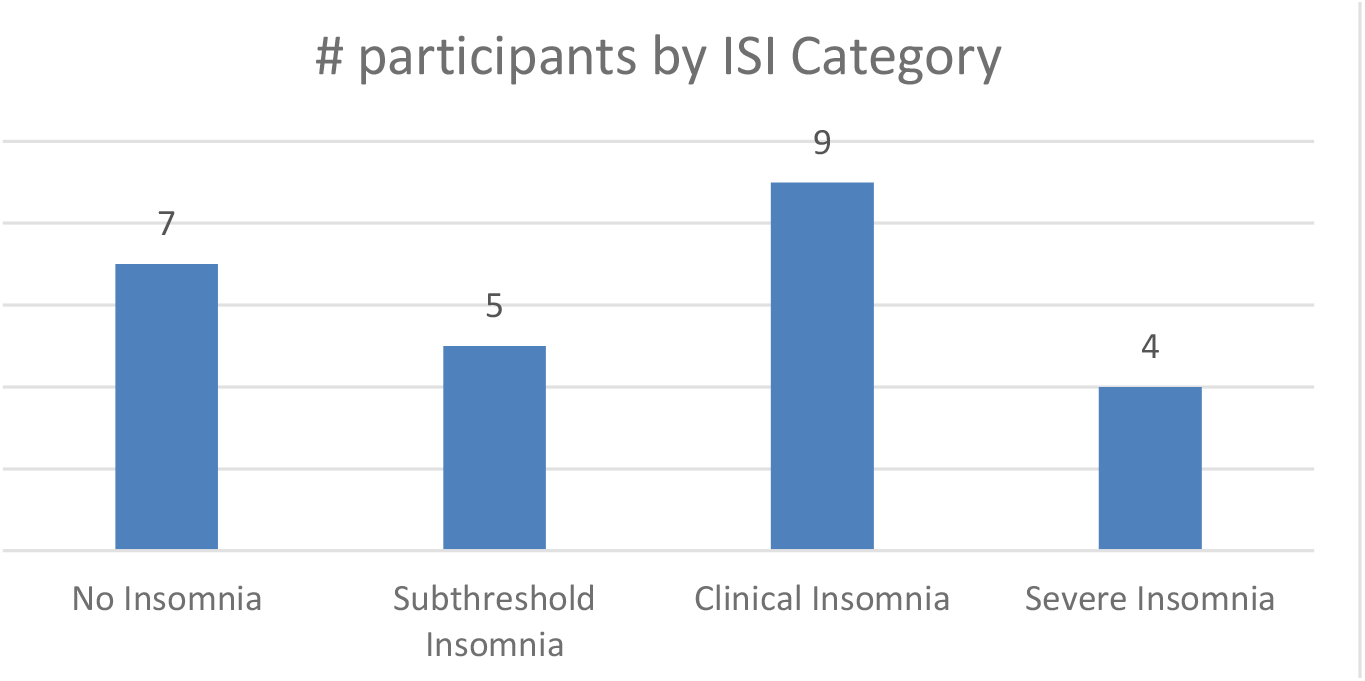
ISI Distribution of participants

For the Fixed tACS condition, a stimulation waveform composed of two sinusoids was created, one at 10Hz (targeting the alpha band) and one at 5Hz (targeting the theta band). The two components were started in phase and thus summed to a maximal possible amplitude with every other cycle of the 10Hz signal. These two frequency bands were targeted given their association with sleep onset, with alpha band activity (8-13Hz) linked to stage I sleep and theta band activity (4-8 Hz) linked to the transition to stage II sleep ^[9]^. The amplitude of both sinusoids was set at 0.6mA intensity (peak-to-peak).

For the Personalized tACS condition, a preliminary session was conducted to collect EEG data from the participants to identify, on an individual basis, the power peaks within the alpha and theta bands. These data were obtained during a 15-minute daytime session, with the participant in a relaxed, eyes-closed state, before any of the pre-sleep stimulation sessions. The custom stimulator device had 2 channels of EEG with electrodes at frontal-lobe sites of Fp1 and Fp2 using a bipotential reference electrode at Fpz.

The EEG signal was band pass filtered with cutoff frequencies at 0.3Hz and 45 Hz. The power spectral density (PSD) was calculated using the Welch method on the filtered data and the Fooof algorithm ^[18]^ was used to determine frequency peaks. A k-means algorithm was used to calculate all peaks between 3-12Hz.

Two peaks were identified from these peaks, the first selected as the peak closest to 5Hz within 4-6Hz band and second selected as the peak closest to 10Hz within 9-11Hz band. The stimulation waveform for the personalized condition was created by combining the sinusoids at the identified two peak frequencies. As with the Fixed stimulation protocol, the amplitude of both sinusoids was set to 0.6mA (peak-to-peak). The two sinusoids in the Personalized stimulation protocol were started in phase, but they were not harmonics.

Each participant used the device over a micro-longitudinal 2-week intervention period. For one of the weeks, the app was set to deliver Fixed stimulation for 15 minutes. For the other week, the app was set to deliver Personalized stimulation for 15 minutes. Participants were asked to use the headset “as often as convenient”, with the recognition that they were unlikely to use the device on each night. Days in which the participants did not wear the headset (no stimulation) or when they failed to use the app to administer stimulation served as a Control condition (with data available only if they were wearing the tracker device). On average, Fixed stimulation was administered on 5 days (range: 3-6), Personalized on 4 days (range: 3-6), and 3 days of Control data was obtained (range: 1-6).

### Tables and Charts: Included at the end of the manuscript

## Data Analysis

Sleep tracking data was obtained using a Fitbit tracker, the output of which provided sleep/wake durations for the participant through the night. Sleep stage data from the device were not analyzed since the classifier accuracy of different sleep stages for the Fitbit tracker are not sufficiently robust. On days with Fixed or Personalized tACS, sleep onset was defined as the interval between the end of stimulation and the start of the first sleep epoch as determined by the tracker data.

As stimulation sensation was noticeable for the participant, we used the end of stimulation as the event from which to begin measuring sleep onset. Given that there was no stimulation in the Control condition, sleep onset data (as defined to be measured from the end of stimulation) was not recordable for this condition.

Data outliers were defined based on two pre-hoc criteria: 1) If the sleep start time for a session was beyond ±1.5 hour of their sleep start time distribution inter-quartile-range, or 2) the participant’s sleep duration fell beyond ±1.5 hour of their sleep duration distribution inter-quartile-range. On average, 0.3 sessions (range: 0-2) were excluded and the distribution was similar across the conditions. From the remaining data, mean sleep onset and sleep duration scores for each participant were calculated in each condition. Note that the Control data were collected across the two-week study period.

## Results

### Personalized vs. Fixed tACS Stimulation

Personalized tACS stimulation increased sleep duration by 22 minutes compared to the Control condition (p=.04) and 19 minutes compared to Fixed stimulation (p=.03; see Figure 3). Fixed stimulation increased sleep duration by 3 minutes compared to Control condition, but this difference was not significant (mean: 3 minutes; p=0.75). Personalized stimulation resulted in a faster sleep onset by 6 minutes compared to the Fixed stimulation (28% improvement, p=.02) – refer to Figure 4, Table 1 and Table 2.

**Figure 3:**
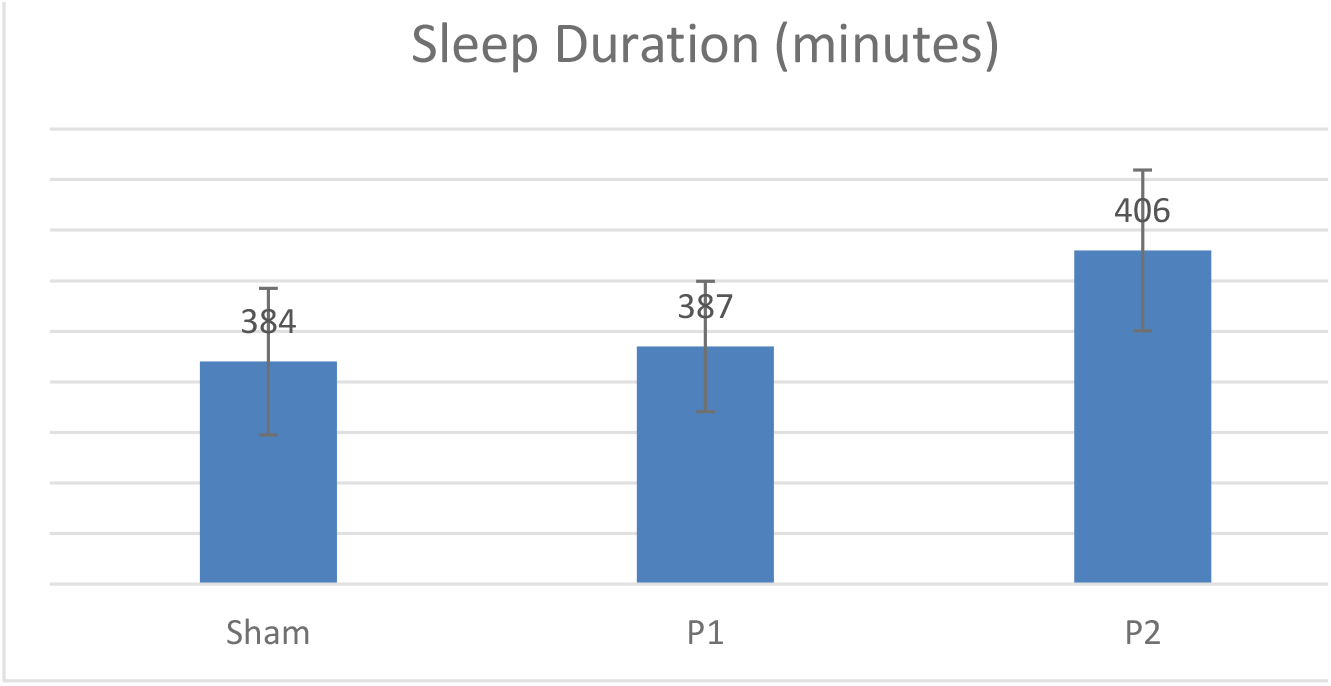
Sleep Duration comparison across conditions

**Figure 4.**
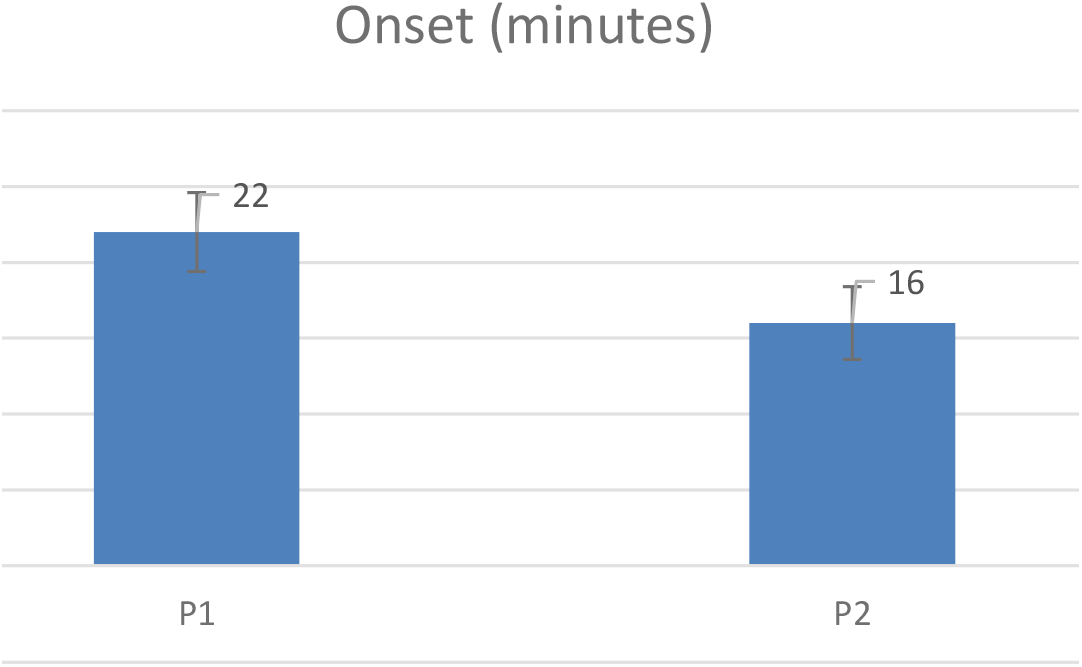
Onset Comparison across stimulations

**Table 1.**
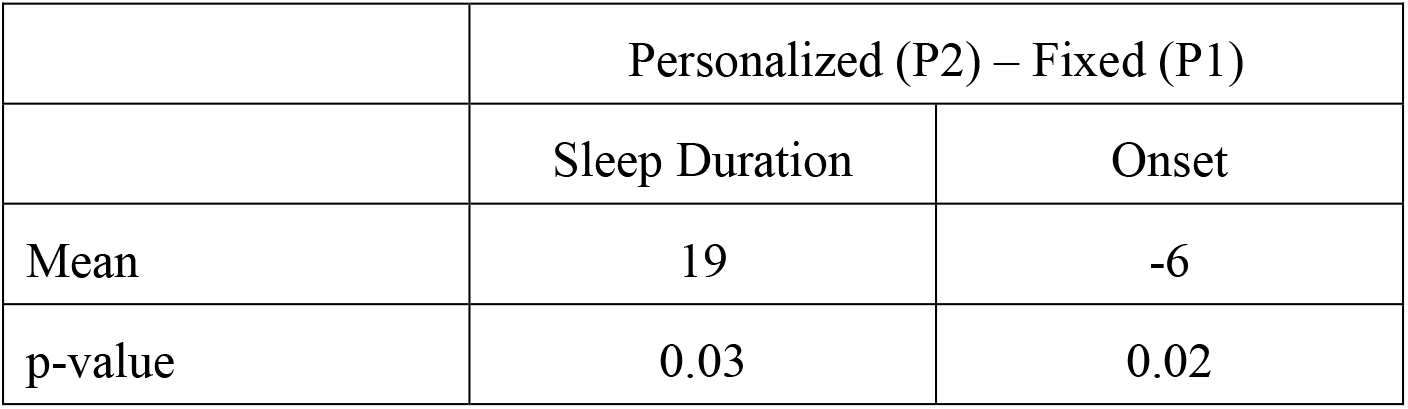
Sleep Duration and Onset Performance

|  | Personalized (P2) – Fixed (P1) |  |
| --- | --- | --- |
|  | Sleep Duration | Onset |
| Mean | 19 | -6 |
| p-value | 0.03 | 0.02 |

**Table 2.** Sleep Duration compared to sham

|  | Sleep Duration Compared to Sham |  |
| --- | --- | --- |
|  | Fixed (P1) | Personalized (P2) |
| Mean | 3 | 22 |
| p-value | 0.75 | 0.04 |

Using the demographic information collected concerning age and sleep hygiene, two secondary analyses were conducted on sleep duration (Table 3). For the age analysis, we divided the participants into two groups based on age: <=50 years old with n=13, >50 years old with n=12.

**Table 3.** Sleep Duration by Age and ISI

|  | Personalized (P2) – Fixed (P1) |  |  |  |
| --- | --- | --- | --- | --- |
|  | Young (<50yrs) | Old (>50yrs) | ISI 1 or 2 | ISI 3 or 4 |
| Mean | 27 | 10 | 4 | 33 |
| p-value | 0.02 | 0.45 | 0.67 | 0.02 |

Age Analysis: First, to confirm the classic age-related decrease in sleep as a validation of the cohort and its normative sleep, there was a significant correlation between age and sleep duration (data from Control condition, r=-0.19, p<0.001), with an average -0.8-minute decrease in sleep duration with every year increase in age (Figure 5), consistent with published norms ^[35]^.

**Figure 5.**
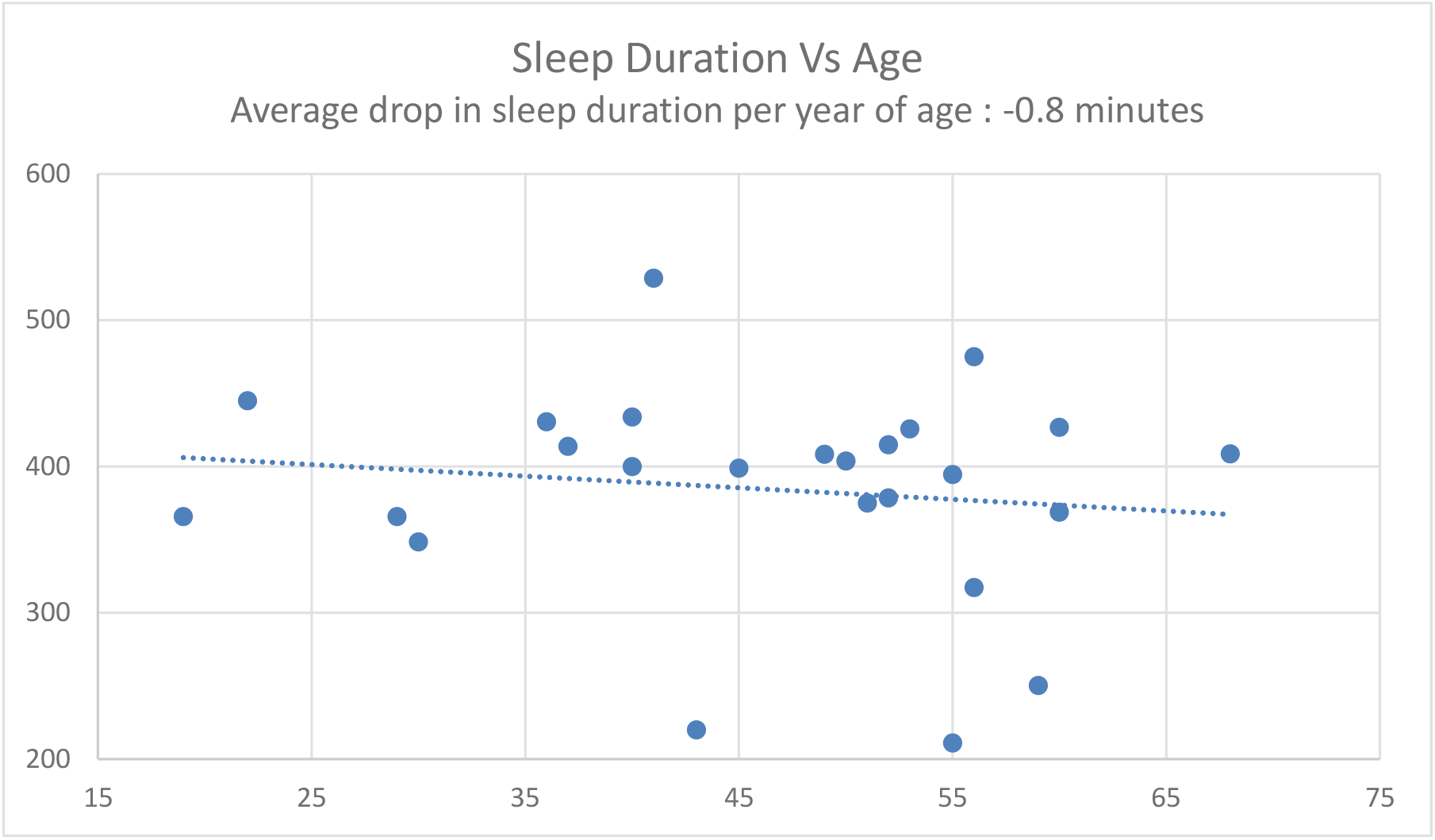
Sleep Duration vs Age Regression

Segmenting the cohort based on age, and first focusing on the younger cohort (<=50 years old), Personalized tACS stimulation resulted in a 27-minute increase in sleep duration, relative to Fixed tACS stimulation (p=0.02), and a 29-minute increase in sleep duration compared to the Control condition (p=0.02). In the older cohort (>50 years old), Personalized tACS stimulation elicited a 10-minute increase in sleep duration, relative to Fixed tACS stimulation (p=0.4) and a 14-minute increase relative to Control condition (p=0.45).

Poor-vs. Good-sleeper Analyses: Based on sleep hygiene, the cohort was further segmented post-hoc into two groups: a normative sleep group (those with no-insomnia and subclinical threshold insomnia ISI categorization, n=12), and a poor sleep group (those with clinical insomnia and severe insomnia ISI categorization, n=13). For the poor sleep group, Personalized tACS stimulation improved sleep duration by 33 minutes compared to Fixed tACS stimulation (p=0.02), and 30 minutes compared to Control condition (p<0.1). For the normative sleep group, Personalized tACS stimulation increased sleep duration by 4 minutes relative to Fixed tACS stimulation (p=0.67; ns) and increased 13 minutes compared to Control condition (p=0.2; ns).

## Discussion

To date, a number of studies have provided evidence that external, non-invasive stimulation of varied forms can enhance sleep quality ^[5] [6] [7] [8]^. For modulating electrical brain activity, these include tDCS, tACS and rTMS, all applied in an open loop i.e., fixed-stimulation manner, and all applied during sleep. Indeed, tACS and rTMS protocols targeting oscillatory patterns in different frequency bands (0.5Hz to 16Hz) using fixed stimulation waveforms across all subjects have led to improvements in several sleep metrics ^[21][39]^.

The current study took a different approach. Specifically, the study tested whether tACS before (rather than during) sleep would similarly improve sleep, and furthermore whether a novel personalized tACS stimulation would further augment sleep effects compared to fixed tACS stimulation.

The results suggest that the tACS stimulation prior to sleep (across the two active conditions) does improve sleep duration, relative to null-stimulation Control condition (12 more minutes of sleep). Furthermore, comparing the two active stimulation conditions, the results suggest that the Personalized tACS stimulation improves sleep duration and sleep onset relative to Fixed tACS stimulation (19 more minutes of sleep and 6 minutes earlier onset of sleep).

An age-related analysis validated the known sleep norms of sleep duration decreasing with age (decrease of 0.8 minutes per year observed with the Control condition). Previous studies have shown that night-to-night and inter-participant variability in sleep quality increases in older cohorts ^[40]^. Given these norms of aging and sleep quality deterioration, we did a post-hoc analysis of the impact of Personalized tACS stimulation on sleep duration relative to Control condition, for a younger cohort (<=50yrs old) and an older cohort (>50yrs old). We observed a robust 29-minute increase in sleep duration for the younger cohort. For the older cohort, sleep duration increased by 14-minutes, providing a potential for using the Personalized tACS stimulation to improve sleep quality with age. However, this increase was not significant and needs further exploration with a larger aging cohort.

Beyond age as a factor, insomnia symptomatology was further analyzed. This was motivated by the factor that current sedative hypnotics have non-trivial side effects, have challenging aspects of long-term efficacy, may fail to implement normative sleep physiology ^[2]^, the recent American College of Physicians recommendation that they should no longer be a first line treatment approach for those with sleeping difficulties ^[2]^.

Post-hoc analysis demonstrated that the poor sleep sub-group (defined as having significant insomnia categorization using the ISI) showed a larger boost in sleep duration (33 minutes increase) from personalized tACS stimulation compared to the Fixed tACS protocol. As a point of contrast, the typical prescription sleep medication, zolpidem (brand name, Ambien), has been shown to increase total sleep time by 35.5 minutes relative to placebo ^[36]^, and a more recent medication, suvorexant (brand name, Belsomra), has a reported increase in total sleep time of 28 minutes ^[37]^. As such, the current results offer early tentative evidence that non-invasive, personalized stimulation may be a future viable alternative intervention for insomnia, should such finding be replicated large-scale.

Finally, the current study administered the tACS stimulation for 15-minutes pre-sleep and involved no additional stimulation during sleep. With participants aware of stimulation sensations, this approach has been selected based on multiple studies showing that the benefits of stimulation last well beyond the stimulation period ^[10][11][29]^. That said, we can see two important extensions of the current approach in future studies. First, resting state EEG could be obtained prior to each night to account for within-subject fluctuations across days in peak frequency. Second, EEG could be monitored during sleep to allow for additional closed-loop stimulation during the night.

### Limitations

The sample size of 25 in the present study is modest, especially when considering the large age range, variance in ISI scores, and the fact that participants self-administered tACS stimulation in their home setting at a time of night that was variable. Clearly this work needs to be replicated and extended in studies with samples of significantly greater size, and within targeted subgroups (e.g., age, sleep hygiene).

In the Fixed tACS stimulation, the two frequencies (5Hz and 10Hz) are harmonics. Thus, this condition differs from the Personalized in the phase or temporal coherence of the composite waveform given that there will be a synchronized maximum every 2 cycles of the 10Hz. This pattern is not present in the Personalized tACS stimulation. Given that the Fixed condition failed to produce a benefit, it may be that having a consistent phase relationship offsets the effects of the base frequencies. This issue can be addressed in future work by using a fixed pattern in which the two components are not harmonics. It would also be useful to compare two individualized stimulation patterns assigned to each individual, one based on their own EEG recordings and a control condition involving another individual’s EEG recordings.

The null-stimulation Control condition in this current study did not represent a classical sham stimulation (which typically involves a short ramp-up period of stimulation lasting approximately 15 seconds and ramp-down). While this is an experimental weakness, it is important to recognize that participants are unlikely to be fully blinded with our current mode of stimulation; in general, participants were aware of sensation when the stimulation was on. Nonetheless, a typical sham control should be implemented in future studies.

Given previous results using tACS in the alpha and theta range, we were surprised that the Fixed tACS stimulation failed to produce a significant improvement on the sleep measures compared to the Control condition. This null result could have resulted from a lack of power in terms of sample size within the sub-cohorts or some of the other factors discussed above (e.g., use of a composite involving harmonics). Nonetheless, the potential of non-invasive brain stimulation is supported by our finding that personalized tACS stimulation provided significant improvements relative to fixed tACS stimulation and the control condition.

## Conclusions

These findings provide evidence that frequency specific personalized electrical brain stimulation offers a promising method for optimizing human sleep quality. Moreover, these effects were especially prominent in those with the highest ratings of insomnia, suggesting a potential future therapeutic opportunity, once replication of these findings has been accomplished in larger cohorts.

## Acknowledgments

We would like to acknowledge Aaron Bromberg, the CEO of StimScience Inc. for his support helping organize and conduct this study. We also would like to acknowledge Roslyn Krishna, Jeff Fernandez, and Sebastian Mirano for their contributions in the execution of the study.

